# Lake size shapes the relationship between body mass and gut microbiota in threespine stickleback (*Gasterosteus aculeatus*)

**DOI:** 10.1101/2025.12.01.691584

**Authors:** Sihan Bu, Sania Chaudhary, Rebecca Kramer-Earley, Kelly Ireland, Jolie Atwood, Daniel I. Bolnick, Andrew P. Hendry, Catherine L. Peichel, Natalie C. Steinel, Jesse N. Weber, Grant E. Haines, Alison M. Derry, Kathryn Milligan-McClellan

**Affiliations:** Department of Molecular and Cell Biology, University of Connecticut, Storrs, CT, USA; Department of Biology, University of Alaska Anchorage, Anchorage, AK, USA; Department of Biology and Wildlife, University of Alaska Fairbanks, Fairbanks, AK, USA; Department of Ecology and Evolutionary Biology, University of Connecticut, Storrs, CT, USA; Department of Biology, McGill University, Montréal, Québec, Canada; Division of Evolutionary Ecology, Institute of Ecology and Evolution, University of Bern, Bern, Switzerland; Department of Biological Sciences, University of Massachusetts, Lowell, MA, USA; Center for Pathogen Research and Training, University of Massachusetts, Lowell, MA, USA; Department of Integrative Biology, University of Wisconsin-Madison, Madison, WI, USA; Aquaculture and Fish Biology, Hólar University College, Sauðárkrókur, Iceland; Sciences Biologiques, Université du Québec á Montréal, Montréal, Québec, Canada

**Keywords:** threespine stickleback, *Gasterosteus aculeatus*, gut microbiota, lake size, mass

## Abstract

Host-microbe interactions are shaped by both host and environmental factors. However, little is known about how host-microbe interactions vary across populations within a species. Here, we characterized the gut microbiota of 191 wild threespine stickleback fish (*Gasterosteus aculeatus*) from six populations from Alaskan lakes spanning a gradient of surface area. We tested how environmental context (lake size and ecotype) and host traits (sex, body mass, gravidity, *Schistocephalus solidus* (*S. solidus*) infection, and fibrosis) influence stickleback gut microbial composition using 16S rRNA gene sequencing. We found that the lake surface area strongly predicted fish gut microbial alpha diversity. Fish from intermediate-sized lakes harbored significantly more diverse microbiota than those from small and large lakes, independent of ecotype. Body mass was associated with gut microbial diversity. Model-predicted marginal effects from the mass and lake surface area interaction analysis showed that the association between fish mass and microbial alpha diversity was strongly negative in the smallest lakes, weakest in intermediate-sized lakes, and strongly positive in the largest lakes. In addition, sex and *S. solidus* infection were significantly associated with gut microbiota alpha and beta diversity, whereas fibrosis and gravidity showed minimal effects. Differential abundance analysis revealed lake size-dependent associations between body mass and individual taxa. Together, these results demonstrate that both habitat context and host variation interactively shape stickleback gut microbial communities in the wild. Integrating lake-level and individual-level analyses reveals how ecological setting modulates host-microbe associations, offering insights into the role of the gut microbiota in host adaptation and population divergence.

## Introduction

The gut microbiota, a complex community of bacteria, archaea, viruses, and eukaryotes residing in the gastrointestinal tract, plays a crucial role in the biology, ecology, and evolution of hosts [1, 2]. Over host evolution, symbiotic gut microbial communities contribute to host adaptations to different habitats by regulating essential metabolic functions, immunity, digestion, and more [3]. The threespine stickleback fish (*Gasterosteus aculeatus*) is an established model organism for gut microbiome studies because it occupies diverse habitats and exhibits pronounced ecological and morphological variation. Stickleback populations span marine to freshwater systems and commonly diverge into benthic and limnetic ecotypes within freshwater [4, 5]. This diversity makes stickleback an ideal system for investigating how host traits and environmental context shape gut microbial communities. Benthic sticklebacks typically evolve deeper bodies and wider mouths, feeding on invertebrates in the sediment, while limnetic forms develop smaller bodies suited for open-water zooplankton feeding [4, 6, 7]. Previous studies showed that stickleback gut microbiota differed between ecotypes, reflected population-specific diets, and varied with host genetic background and environmental context in both wild and lab populations [8–11]. However, many lakes only contain one stickleback ecotype, and thus it often covaries with other environmental factors in a particular lake [12]. As a result, lake-level factors such as lake surface area may be important yet underexplored drivers of fish gut microbial communities. Studies on sticklebacks have suggested that intermediate-sized lakes supported high habitat and diet heterogeneity, leading to higher among-individual diet variation compared to small and large lakes [13]. Indeed, a recent study on stickleback revealed that greater gut microbiota uniqueness was observed in individuals from intermediate-sized lakes, indicating that diet diversity may promote gut microbiota diversity [14]. These observations suggest that lake surface area may play a crucial role in structuring gut microbiota and can also influence the relationships between host traits and microbial diversity.

Several host traits, including body mass, parasite infection, immune responses, sex, and reproductive status, have been also investigated in shaping the gut microbiota of stickleback and other fish species. Body mass may influence gut microbiota through effects on feeding behavior, habitat access, gut morphology, and microbial colonization potential [15, 16]. In *Gymnocypris chilianensis*, heavier individuals showed significantly greater gut microbial richness and diversity [17]. Because lake surface area can restructure resource availability and ecological opportunity, it may also modulate the relationship between fish body mass and gut microbial diversity. However, whether the influence of body mass on stickleback gut microbiota varies across different lake contexts remains untested. Additionally, the tapeworm *Schistocephalus solidus* (*S. solidus*) alters stickleback gut microbial communities and host immune system, indicating a potential indirect pathway by which parasite-mediated immune modulation shapes the gut microbiome[18–22]. Weighted and unweighted UniFrac distances of beta diversity showed significant separation between *S. solidus*-infected and non-infected wild Alaskan stickleback populations [23]. Similarly, fibrosis represents an immune response to *S. solidus* infection in stickleback and may also influence the intestinal environment, thereby influencing gut microbial composition, though direct tests remain limited [24, 25]. Female fish tended to have higher Amplicon Sequence Variant (ASV) richness than male fish among wild stickleback populations [10]. In sticklebacks, female reproductive effort (clutch size relative to body size) exhibits plasticity, suggesting that gravidity may modulate energy allocation in ways that alter gut microbial communities [26]. Together, these factors motivate our focus on how interactions among environmental context and host traits jointly influence stickleback gut microbial composition.

To address these gaps, we investigated the gut microbiota of 191 threespine stickleback collected from six Alaskan lakes. We tested how environmental factors (lake surface area and ecotype) and host traits (sex, mass, gravidity, *S. solidus* infection, and fibrosis) shape stickleback gut microbial composition. Additionally, we investigated whether lake size affected the relationship between body mass and gut microbiota and conducted differential abundance analyses to identify taxa associated with this relationship. This study explicitly links habitat variation and host traits to fish gut microbiota structure across ecological scales. By combining lake-level environmental factors with individual-level host data, we provide a framework to highlight the importance of integrating environmental and host-level data to better understand the assembly of gut microbiota in natural ecosystems.

## Methods

### Fish sample collection

Fish were collected in a quarter inch mesh unbaited minnow traps set overnight on the shoreline of four lakes in the Matanuska-Susitna (Mat-Su) Valley and four lakes in the Kenai Peninsula Alaska in June 2019 as part of an experiment described in a previous paper [27]. Gut samples were initially collected from eight lakes; however, two lakes were excluded from subsequent analyses because sample labels were lost during processing, which prevented accurate identification. We analyzed 40 fish per lake with a total of six lakes for subsequent analyses (Table 1). Lake properties and ecotype assignments of the fish (i.e. benthic or limnetic) are as previously described [27]. Briefly, fish were euthanized with buffered MS-222, then weighed and measured for standard length. The sex of each fish was determined by visual inspection of gonad morphology. Fibrosis in the peritoneal cavity was scored on a scale from 0 to 4, where 0 indicated no fibrosis and 4 represented severe fibrosis [28]. *S. solidus* presence was determined by visualization of the fish body cavity. Following these measurements, the entire digestive tracts were dissected using sterile dissecting tools into 1.5mL microcentrifuge tubes that were stored on dry ice. Samples were shipped to the University of Alaska Anchorage and stored at −80°C. Sample collections were done in compliance with Alaska Department of Fish and Game permit P19-19-005 and UAA IACUC protocol 1362929-1.

**Table 1.** The characteristics of the stickleback fish across all the lakes.

| Characteristics | Overall (N=191) | Long Lake (N=34) | Walby Lake (N=35) | South Rolly Lake (N=37) | Tern Lake (N=16) | Spirit Lake (N=39) | Finger Lake (N=30) | p-Value |
| --- | --- | --- | --- | --- | --- | --- | --- | --- |
| Sex, n(%)<sup>1</sup> |  |  |  |  |  |  |  |  |
| Male | 48 (25.13%) | 14 (41.18%) | 8 (22.86%) | 8 (21.62%) | 3 (18.75%) | 9 (23.08%) | 6 (20%) | 0.31 |
| Female | 142 (74.35%) | 19 (55.88%) | 27 (77.14%) | 29 (78.38%) | 13 (81.25%) | 30 (76.92%) | 24 (80%) |  |
| Gravidity (Females Only), n(%) |  |  |  |  |  |  |  |  |
| Gravid | 44 (30.99%) | 1 (5.26%) | 8 (29.63%) | 4 (13.79%) | 6 (46.15%) | 16 (53.33%) | 9 (37.5%) | <0.001 |
| Non-gravid | 98 (69.01%) | 18 (94.74%) | 19 (70.37%) | 25 (86.21%) | 7 (53.85%) | 14 (46.67%) | 15 (62.5%) |  |
| Mass(g), median (IQR) | 1.16 (0.66) | 1.23 (0.78) | 1.09 (0.36) | 0.88 (0.46) | 1.53 (1.18) | 1.2 (0.9) | 1.43 (0.48) | <0.001 |
| Length(mm), median (IQR) | 46.4 (9.32) | 46.79 (12.51) | 43.62 (4.36) | 47.26 (5.87) | 48.62 (14.26) | 45.54 (12.24) | 49.1 (4.85) | 0.001 |
| *S. solidus* infection, n(%) |  |  |  |  |  |  |  |  |
| Infected | 21 (11%) | 0 | 17 (48.57%) | 0 | 2 (12.5%) | 0 | 2 (6.67%) | <0.001 |
| Non-infected | 170 (89%) | 34 (100%) | 18 (51.43%) | 37 (100%) | 14 (87.5%) | 39 (100%) | 28 (93.33%) |  |
| Fibrosis score, n(%) |  |  |  |  |  |  |  |  |
| 0 | 135 (70.68%) | 22 (64.71%) | 32 (91.43%) | 33 (89.19%) | 12 (75%) | 18 (46.15%) | 18 (60%) | <0.001 |
| 1 | 25 (13.09%) | 5 (14.71%) | 3 (8.57%) | 1 (2.7%) | 3 (18.75%) | 8 (20.51%) | 5 (16.67%) |  |
| 2 | 26 (13.61%) | 5 (14.71%) | 0 (0%) | 3 (8.11%) | 1 (6.25%) | 10 (25.64%) | 7 (23.33%) |  |
| 3 | 5 (2.62%) | 2 (5.88%) | 0 (0%) | 0 (0%) | 0 (0%) | 3 (7.69%) | 0 (0%) |  |
| Ecotype |  | Limnetic | Benthic | Limnetic | Benthic | Limnetic | Benthic |  |
| Lake surface area (ha) |  | 16.9 | 19.6 | 43.8 | 44.1 | 123.2 | 135.7 |  |
| Geographical region |  | Mat-Su | Mat-Su | Mat-Su | Kenai | Kenai | Mat-Su |  |

<sup>1</sup>Missing information for one sample in Long Lake. Lakes are arranged from the smallest to the largest in terms of surface area.

### DNA extraction and 16S rRNA gene PCR amplification

Frozen guts were thawed for less than 5 minutes at 85°C using heating blocks. Enzymatic lysis buffer (ELB) was prepared according to the Pretreatment for Gram-positive bacteria protocol per DNeasy® Blood & Tissue Handbook (Qiagen, Germany). A total of 300μL ELB and entire fish guts were transferred to Lysing Matric E tubes (MP Biomedicals, USA) and homogenized using a FastPrep-24™ classic bead beating grinder and lysis system (MP Biomedicals, USA) at 6,500rpm for 40 seconds, repeated three times. Bead tubes were placed on ice for at least 1 minute between bead-beating steps. Following homogenization, 200µL of the lysate was transferred to collection microtubes (provided with Qiagen DNeasy® 96 Blood & Tissue kit) after a brief centrifuge step. Samples were incubated at 37°C for 30 minutes. DNA extraction was performed according to the manufacturer’s protocol with a few modifications. Specifically, a working solution was prepared by combining 25µL of Proteinase K and 200µL of Buffer AL (without ethanol) per sample. 225µL of this solution was added to each collection microtube, followed by incubation at 56°C for 30 minutes. The tubes were then covered and vigorously shaken for 15 seconds. 20µL of ethanol was added, and the samples were thoroughly mixed by vortexing. The protocol was then continued from Step 7 of the manufacturer’s instructions (October 2022). DNA was quantified using a Qubit 4 Fluorometer (Thermo Fisher Scientific, USA).

Library preparation and sequencing were performed at the University of Connecticut Microbial Analysis, Resources, and Services (MARS) facility. Briefly, the V4 region of the 16S rRNA was amplified using 515F (GTGYCAGCMGCCGCGGTAA) and 806R (GGACTACNVGGGTWTCTAAT) primers from Earth Microbiome Project (EMP) [29, 30] with customized Illumina adapters and 8-base pair dual indices [31]. A total of 15µL PCR mixture including 30ng of DNA, 0.6μL of each forward and reverse primers, 0.15μL Go-Taq DNA polymerase (Promega, USA), 3.3μg Bovine Serum Albumin (BSA) (New England BioLabs, USA) was used. To overcome inhibition from host DNA, 0.1pmol primer without the indexes or adapters was added to the PCR reaction. Reactions were performed in triplicate and amplified using the following thermocycler protocol: 95°C for 2 minutes, 30 cycles of 30s at 95°C, 30s at 50°C, and 60s at 72°C, followed by final extension at 72°C for 10 minutes. PCR amplicons were pooled for quantification and visualization using the QIAxcel DNA Fast Analysis (Qiagen, Germany). PCR products were normalized based on DNA concentration within the 250–400 bp range and pooled using the epMotion 3075 liquid handling robot (Eppendorf, Germany). The pooled products were purified using Mag-Bind® RxnPure Plus kit (Omega Bio-tek, USA) following the manufacturer’s protocol, using a 0.8×bead-to-PCR product ratio. The cleaned 16S library was sequenced on the MiSeq platform with a pair-end v2 2×250 base-pair kit (Illumina, USA).

### Processing and analysis of sequence data

Demultiplexed sequence reads were processed through QIIME2 (version 2023.9) [32]. Sequences with low quality (<30 quality score) were trimmed and truncated using the DADA2 plugin [33]. ASVs were assigned using a pre-trained Naïve Bayes classifier on a full-length 16S rRNA gene SILVA v138 99% OTUs database with feature-classifier plugin [34–36]. Sequences of mitochondria, chloroplasts, archaea, and eukaryota were filtered out before further analysis. Sequencing depth was set at 3,000 reads per sample based on the alpha rarefaction plot to optimize diversity capture while retaining most samples, leaving 191 samples with 1770 ASVs. All the analyses were conducted at the genus taxonomic level.

### Statistical analysis

Data were analyzed using R (version 4.2.2). The differences in sex, gravidity, and *S. solidus* infection among lakes were tested using a binomial general linear model (GLM). Model significance was evaluated by analysis of deviance with χ^2^ tests. Analysis of variance (ANOVA) was used for testing the differences in mass and length among lakes, and Tukey’s HSD was used for post-hoc pairwise comparisons. Spearman’s correlation was used to assess the relationship between mass and length. Differences in fibrosis severity among lakes were tested using the Kruskal-Wallis test.

Overall, gut microbiota alpha and beta diversity were analyzed using the vegan package [37]. For lake-level analyses, we investigated the relationships between log lake surface area, fish ecotype, and gut microbiota composition. The mean alpha diversity (Chao1 and Shannon indices) scores for each lake were calculated and used as the representative value. A quadratic regression model was used to test whether the mean of Chao1 and Shannon scores in each lake had a significant U-shaped relationship with the log lake surface area. ANOVA was used to compare the linear and quadratic regressions to determine if the addition of quadratic terms improved the model fit. The mean scores were also compared between ecotypes (benthic and limnetic) using the Wilcoxon rank-sum test. When analyzing beta diversity, count data were first converted to relative abundance of taxa for each fish, then averaged within each lake to obtain a single community composition per lake. Sorensen and Bray-Curtis dissimilarity metrics were calculated and tested with log surface area and ecotype using Permutational Multivariate Analysis of Variance (PERMANOVA) with the adonis2 function.

For the individual-level analyses, we evaluated whether gut microbial diversity is a function of host traits (sex, mass, gravidity, *S. solidus* infection, and fibrosis), lake-of-origin (a fixed effect), and interactions between lake and individual fish traits. Model selection was guided by Type II sums of squares ANOVA (Supplementary Table 1) and Akaike Information Criterion (AIC) (Supplementary Table 2) [38]. The final model for alpha diversity analyses included lake-of-origin, sex, mass, fibrosis score, and *S. solidus* infection as main effects, together with the interaction terms including lake:mass, sex:fibrosis, sex:*S. solidus* infection, and mass:*S. solidus* infection (Supplementary Table 3). Beta diversity analyses used the same model structure. Because adonis2 cannot test main effects while controlling for interactions, we assessed main effects using marginal tests and interaction terms using sequential tests (Supplementary Table 4-5). To evaluate robustness, we performed two sensitivity analyses. *S. solidus* infection effects were tested in lakes where the parasite was present, and gravidity effects were assessed in female-only individuals. We examined whether lake size influenced the strength of the relationship between fish mass and gut microbiota diversity by testing the interaction between fish mass and log lake surface area on alpha diversity while controlling for the effects of mass, *S. solidus* infection, and lake surface area, as well as the interaction between mass and *S. solidus* infection. We then used the emtrends function from the emmeans package [41] to estimate the marginal effect of mass on each alpha diversity index across the observed range of log lake surface area based on the fitted interaction model.

Associations between fish mass and the relative abundance of individual microbial taxa were tested with Analysis of Compositions of Microbiomes with Bias Correction 2 (ANCOM-BC2) [39] at the genus level on the unrarefied count table. Taxa present in more than 50% of samples were retained for analysis. We included mass, log lake surface area, *S. solidus* infection and sex, along with their interactions (mass:log lake surface area and mass:*S. solidus* infection), based on model selection results, to evaluate whether the relationships between mass and gut microbiota association varied with lake size. Seven taxa showing significant mass and lake surface area interactions (q-value<0.1) were further analyzed in lake-specific models. We then ran separate models within each lake, including *S. solidus* infection as a covariate only in lakes where the parasite was present. P-values were adjusted for multiple comparisons by the Benjamini-Hochberg procedure. All tests were performed with a significance value of 0.05.

## Results

### Characteristics of fish populations and the environment

A total of 191 fish gut samples were included in the final analysis (Table 1). Sex distribution was similar across lakes (2.8% of binomial GLM explained deviance, *p*-value=0.31), with a majority of fish being female (74.35%). Most of the females were not gravid (69.01%). In addition, fish mass differed significantly across lakes (F=5.97, *p*-value<0.001), with South Rolly Lake having the lightest fish (median of 0.88 (interquartile range (IQR)=0.46) grams) when compared to the other five lakes. Fish length was also different between lakes (F=4.3, *p*-value=0.001). However, post-hoc tests revealed no significant differences between any two lakes in terms of fish mass and length. Overall, fish mass was significantly correlated with length (r=0.87, p-value<0.001). *S. solidus* infection prevalence varied significantly among lakes (43.1% of binomial GLM explained deviance, *p*-value<0.001)[40]. Infected individuals were most common in Walby Lake (48.57%) and rare or absent in other lakes. The distribution of fibrosis scores was also significantly different among lakes (χ^2^ =28.36, *p*-value<0.001)[40]. Walby Lake had the lowest prevalence of fibrosis (8.57% of the fish), and Spirit Lake had the highest prevalence of fibrosis (53.85%) (Table 1).

#### Lake surface area was associated with stickleback gut microbiota independently of ecotype

To test the relationship between stickleback fish gut microbiota alpha diversity and lake surface area, we fit quadratic regressions of lake-mean Chao1 and Shannon indices against log lake surface area. For Chao1 index, there was a weak hump-shaped trend on the log lake surface area, but it was not significant (*p*-value=0.2, Figure 1A). However, lake-mean Shannon diversity showed a significant unimodal relationship with log surface area (93% of variation explained, *p*-value=0.009, Figure 1B), indicating that the fish gut microbiota had higher diversity in the intermediate lake surface compared to the ones in small and large lakes. An ANOVA model comparison indicated that adding a quadratic term improved the fit (*p*-value=0.004). For beta diversity analysis, neither fish gut microbiota membership (Sorensen, *p*-value=0.74, Figure 1C) nor composition (Bray-Curtis, *p*-value=0.91, Figure 1D) was associated with log lake surface area. There were no differences in gut microbiota alpha and beta diversity between benthic and limnetic ecotypes (Supplementary Figure 1).

**Figure 1.**
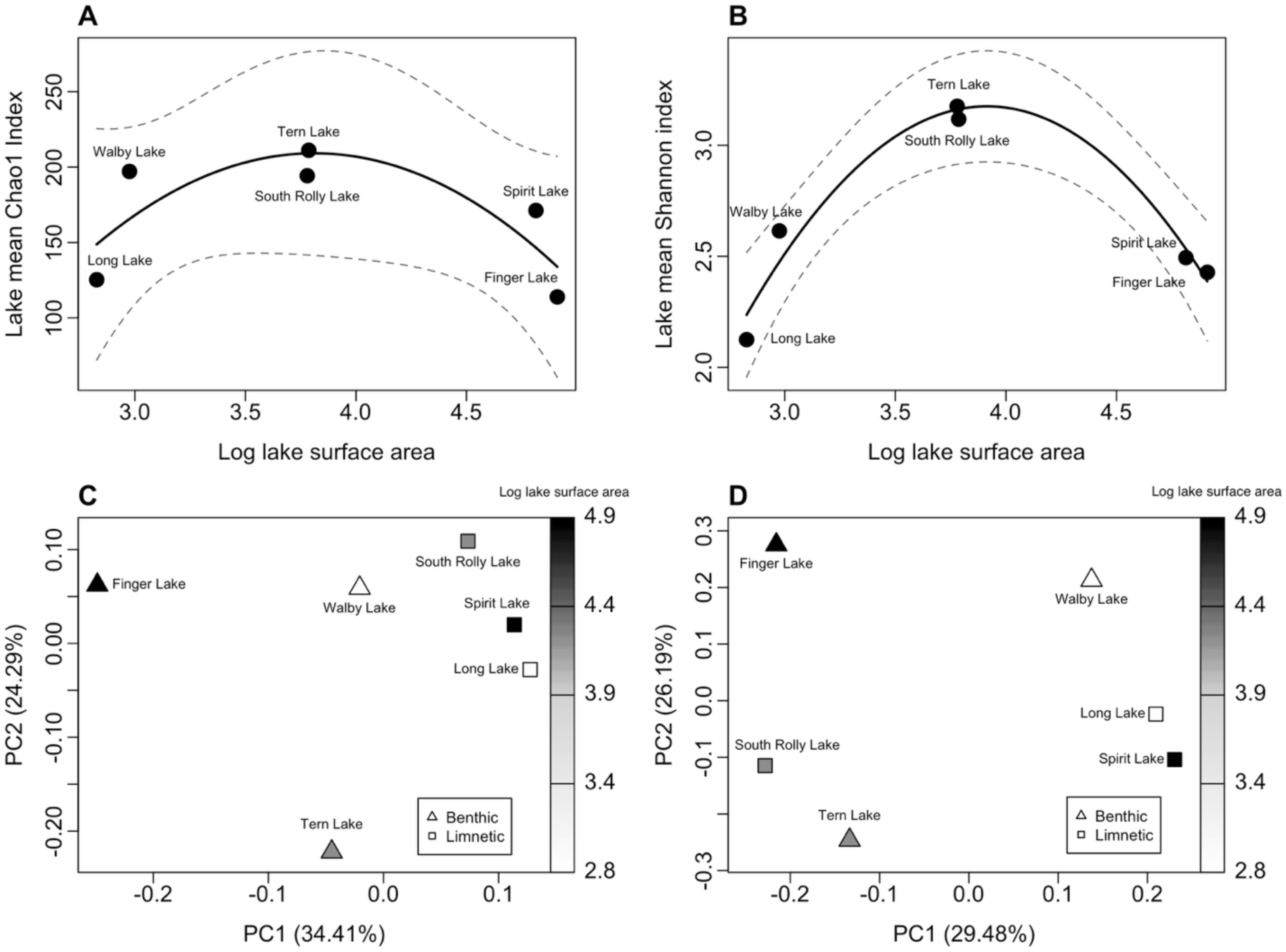
Fish gut microbial composition was associated with lake surface area. For alpha diversity, the quadratic regressions of lake-mean (A) Chao1 (F=2.9, adjusted R^2^=0.43, *p*-value=0.2) and (B) Shannon (F=32.51, adjusted R^2^=0.93, *p*-value=0.009) on log lake surface area were tested, and revealed that stickleback from intermediate-sized lakes had more diverse gut microbiota. Solid lines show quadratic regression curves, and dashed lines indicate 95% confidence intervals. To assess beta diversity, (C) Sorensen (F=0.81, R^2^=0.17, *p*-value=0.74) and (D) Bray–Curtis (F=0.63, R^2^=0.14, *p*-value=0.91) distance metrics revealed that lake size was not associated with fish gut microbial composition. The color gradient shows log surface area. Data points are shaped based on the ecotypes (benthic and limnetic) of the fish.

### Effect of fish mass on gut microbiota composition depended on lake surface area

Fish mass was negatively associated with gut microbiota alpha diversity across lakes (Figure 2). For Chao1 richness, mass had a significant main effect (F=4.95, *p*-value=0.03, Figure 2A), and its interaction with *S*. *solidus* infection was also significant (F=5.34, *p*-value=0.02). A marginally significant interaction between lake-of-origin and mass was observed (F=2.26, *p*-value=0.051), suggesting that the effect of mass on richness may vary among lakes. For Shannon diversity, mass alone did not significantly affect gut microbiota (F=0.91, *p*-value=0.34, Figure 2B); however, the mass and *S. solidus* infection interaction remained significant (F=5.96, *p*-value=0.02). For beta diversity analysis, fish mass did not significantly explain the differences in gut microbiota membership (*p*-value=0.27, Figure 2C) or composition (*p*-value=0.47, Figure 2D). However, a significant interaction between mass and *S. solidus* infection was detected for gut microbiota composition (0.8% variation explained, *p*-value=0.01). These results indicated that the effect of body mass on gut microbiota was context-dependent and potentially shaped by interactions with both parasitic infection and lake environment. Complete statistical results are presented in Supplementary Tables 3-5.

**Figure 2.**
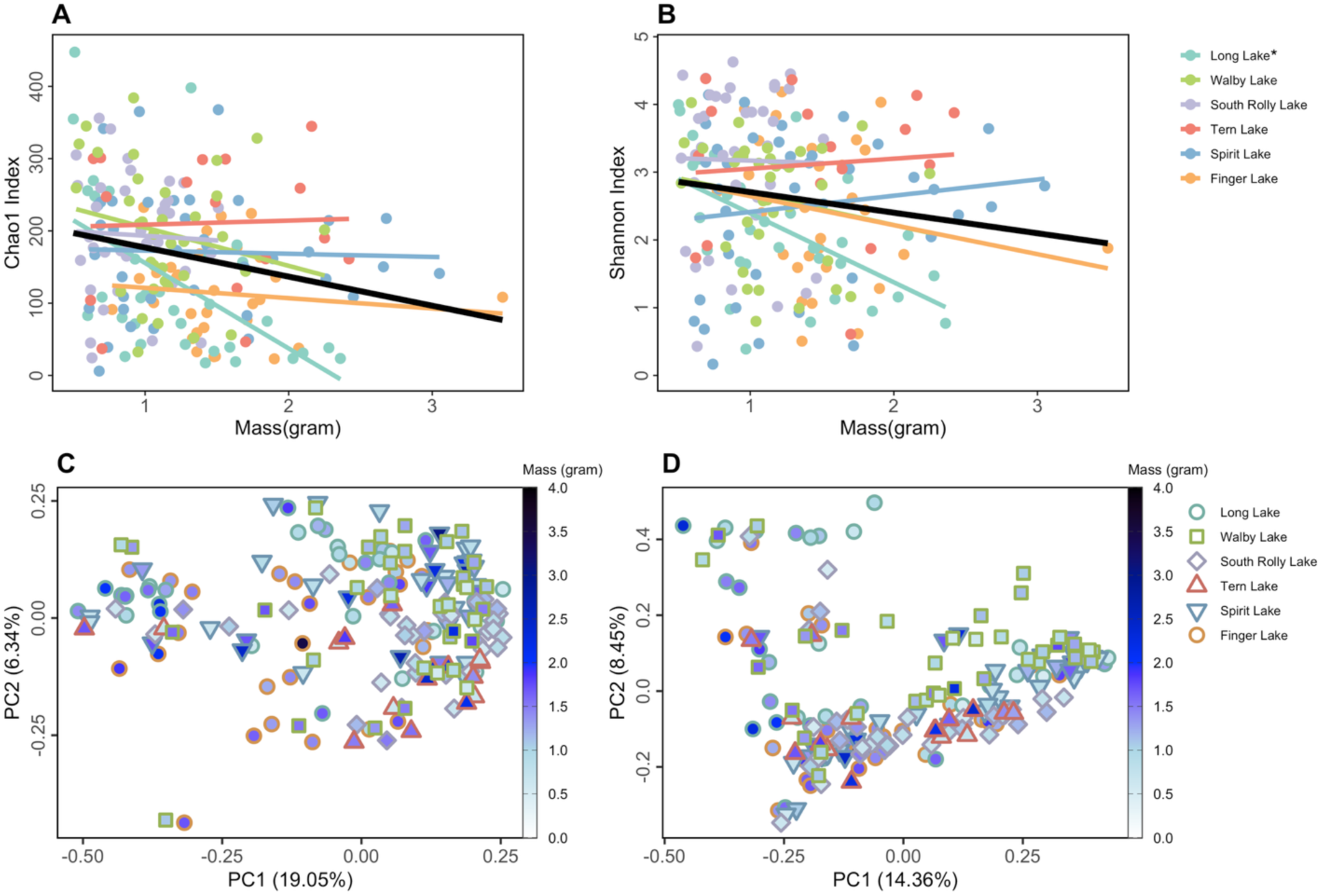
Fish mass was negatively associated with (A) gut microbial richness (Chao1 index, F=4.95, *p*-value=0.03) as measured by alpha diversity. However, the (B) diversity of gut microbiota (Shannon index, F=0.91, *p*-value=0.34) was not associated with fish mass. Fish gut microbiota (C) membership (Sorensen index, F=1.12, R^2^=0.005, *p*-value=0.27) and (D) composition were similar across fish with different mass (Bray-Curtis index, F=0.98, R^2^=0.005, *p*-value=0.47). A total of 190 fish (one sample is missing sex information) were included in the analysis. Colored lines in panels A and B represent lake-specific linear regressions, and the black line indicates the overall relationship for alpha diversity. An asterisk (*) on Long Lake indicates a statistically significant relationship between fish mass and both alpha diversity indices. In panels C and D, colored lines and shapes represent different lakes, and the color gradient within the shape indicates fish mass.

Simple linear regressions fitted within each lake showed that Spirit Lake and Finger Lake (large lakes) exhibited similar trends in the relationship between fish mass and Chao1 richness (Figure 2A). In contrast, Long Lake and Walby Lake (small lakes), as well as South Rolly Lake and Tern Lake (intermediate-sized lakes), displayed comparable but distinct patterns. Because infected stickleback often carry large *S*. *solidus* that can reach a similar weight to their host, higher total mass in infected fish can be driven in part by parasite biomass [41]. However, Walby Lake and Finger Lake contained *S*. *solidus*-infected individuals, whereas Long Lake and Spirit Lake did not. Thus, the observed interaction between mass and *S*. *solidus* infection cannot fully explain the patterns in gut microbiota composition.

Because per-lake analyses revealed substantial variation in the relationship between fish mass and alpha diversity (Figure 2A, Figure 2B), we employed the emtrends function to estimate conditional slopes of fish mass across lake surface areas, which allows the mass effect to vary with lake size instead of assuming a constant slope as in simple linear regression. The interaction between mass and log lake surface area was significant for both Chao1 richness (F=7.57, *p*-value=0.007, Figure 3A) and Shannon diversity (F=5.76, *p*-value=0.02, Figure 3B). The associations between mass and Chao1 richness and Shannon diversity were strongly negative in the two smallest lakes and positive in two largest, but weakest in intermediate-sized lakes (Figure 3). These results indicate that the effect of fish mass on gut microbial richness and evenness depended on lake size, with intermediate-sized lakes exhibiting the weakest relationships between mass and gut microbiota alpha diversity, independent of parasite infection.

**Figure 3.**
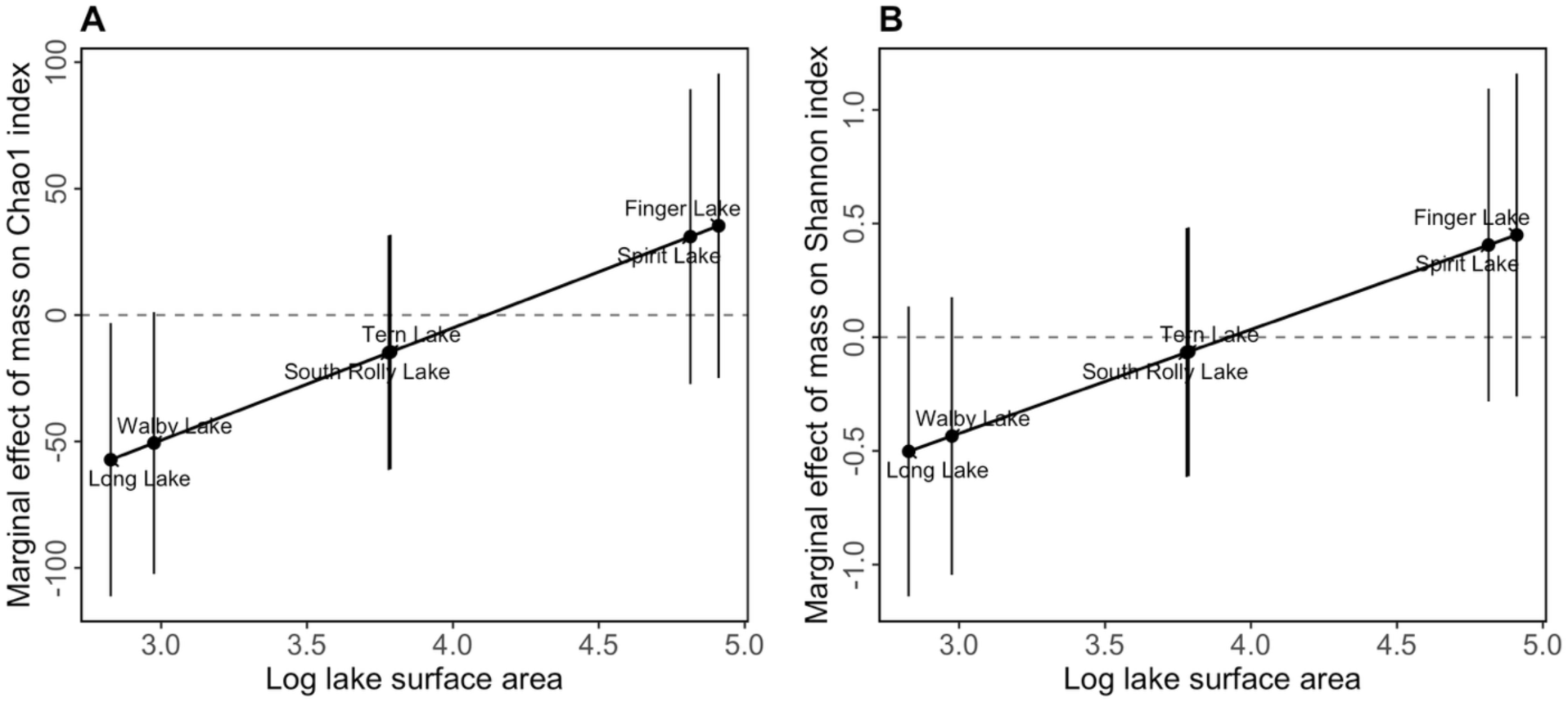
Estimated marginal effects of fish mass on (A) Chao1 index (mass:log surface area interaction: F-value=7.57, *p*-value=0.007) and (B) Shannon index (mass:log surface area interaction: F-value=5.76, *p*-value=0.02), evaluated at each observed log surface area. The estimated effects were the smallest in intermediate-sized lakes compared to small and large lakes. Error bars represent 95% confidence intervals.

### Stickleback gut microbiota composition was associated with sex and S. solidus infection but not fibrosis or gravidity

Female stickleback fish had significantly higher Chao1 (F=4.25, *p*-value=0.04, Figure 4A) and Shannon index (F=7.93, *p*-value=0.005, Figure 4B) than male fish in terms of gut microbial alpha diversity. Male and female fish also differed in microbial membership (2% variation explained, *p*-value<0.001, Figure 4C) and composition (2% variation explained, *p*-value<0.001, Figure 4D) for beta diversity. While no significant interactions between sex and other variables were detected in alpha diversity analysis (Supplementary Tables 3), a significant interaction between sex and fibrosis was observed for Bray-Curtis dissimilarity of beta diversity (2% variation explained, F=1.44, *p*-value=0.01, Supplementary Tables 5). Overall, these results demonstrate that sex is strongly associated with stickleback gut microbiota diversity and community structure, with only limited evidence that this effect is modulated by fibrosis infection.

**Figure 4.**
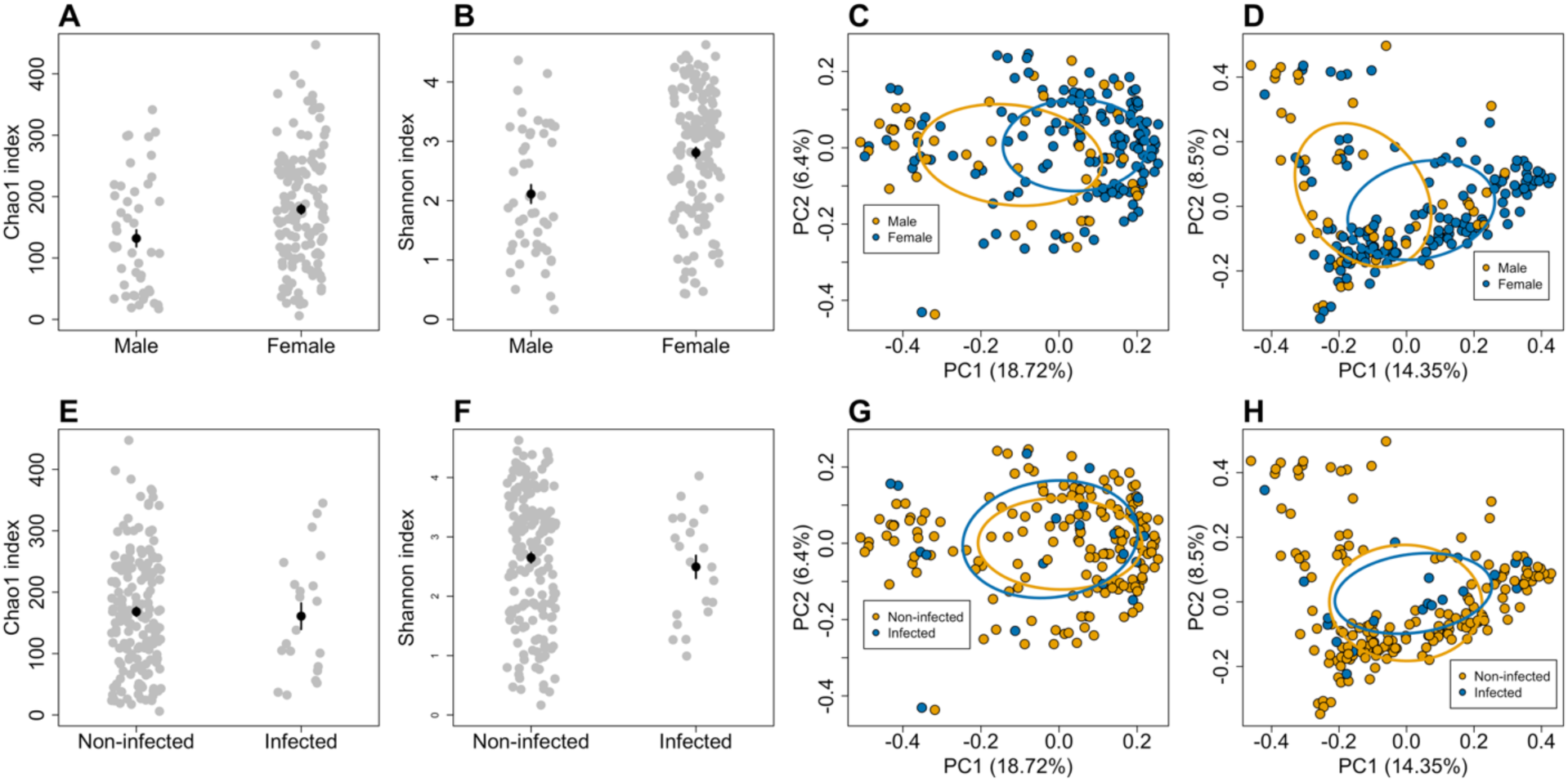
Sex and *S. solidus* infection were associated with stickleback gut microbiota alpha and beta diversity. (A) Females had higher richness of gut microbiota than males as measured by Chao1 (F=4.25, *p*-value=0.04). (B) More diverse gut microbiota were also found in female fish (F=7.93, *p*-value=0.005). For beta diversity, sex influenced gut microbiota (C) membership (F=4.57, R^2^=0.02, *p*-value<0.001) and (D) composition (F=3.56, R^2^=0.02, *p*-value<0.001). (E) *S. solidus* infection was significantly associated with reduced gut microbiota richness (Chao1 index) in stickleback fish (F=4.83, *p*-value=0.03) but not with Shannon diversity (F=1.2, *p*-value=0.28). *S. solidus* infection did not influence gut microbiota community structure based on (G) Sorensen matrix (F=1.57, R^2^=0.007, *p*-value=0.055) or composition measured by (H) Bray-Curtis dissimilarity (F=1.21, R^2^=0.006, *p*-value=0.21). Black dots indicate the mean, and error bars are the mean±SE. Ellipses indicate 95% confidence intervals around centroids.

*S*. *solidus* infection was associated with significantly reduced gut microbial richness as measured by the Chao1 index (F=4.83, *p*-value=0.03, Figure 4E) among all samples. When restricting the analysis to lakes containing *S*. *solidus*-infected fish as a sensitivity test, the direct association between *S*. *solidus* infection and Chao1 richness was not significant, though a trend was observed (F=3.39, *p*-value =0.07, Supplementary Table 3). No significant effect of *S. solidus* infection on Shannon diversity was observed in either the sensitivity analysis or full dataset. The interaction between mass and S. *solidus* infection remained significant on Chao1 (F=4.13, *p*-value=0.046) and Shannon indices (F=7.91, *p*-value=0.006) in sensitivity analysis, consistent with results from the full dataset. This indicates that the association between fish mass and gut microbial diversity was highly dependent on *S*. *solidus* infection status. In addition, a significant interaction between sex and *S. solidus* infection (F=4.33, *p*-value=0.041) was detected for Shannon diversity in sensitivity analysis, suggesting that the impact of infection on gut microbiota diversity may differ between males and females. Analyses of beta diversity showed no significant impact of *S. solidus* infection on microbial community structure using Sorensen dissimilarity or on community composition using Bray-Curtis dissimilarity across all lakes and within the subset of lakes where *S*. *solidus* infection was present (Figure 4G-4H, Supplementary Table 4). However, the interaction between mass and *S*. *solidus* infection had a significant effect on gut microbiota composition when measured with Bray-Curtis dissimilarity (all-lake analysis: F=1.84, R² =0.008, *p*-value=0.01; sensitivity analysis: F=2.1, R² =0.02, *p*-value=0.005), even though *S*. *solidus* infection alone was not significant. Together, these findings conclude that fish mass and *S*. *solidus* infection each had significant effects on gut microbiota composition, with an additional interaction indicating that the influence of one trait varied depending on the other. Complete statistical results are presented in Supplementary Tables 3-5.

Fibrosis did not affect fish gut microbiota alpha or beta diversity (Supplementary Figure 2A-D). Similarly, among female fish, gravidity alone did not affect alpha or beta diversity (Supplementary Figure 2E-H). Gravidity showed a trend toward influencing gut microbiota composition when measured with Bray-Curtis dissimilarity (F=1.46, R²=0.009, *p*-value=0.07; Supplementary Figure 2H). In contrast, the interaction between gravidity and *S*. *solidus* infection significantly influenced gut microbiota composition (F=1.65, R²=0.01, *p*-value=0.03; Supplementary Table 5).

### Differential abundance analysis identified lake size-dependent effects of fish mass on gut microbiota

Given that the association between fish mass and gut microbiota diversity depended on lake size, we next investigated whether specific microbial taxa underlay these community-level patterns. To this end, we performed differential abundance analysis using the ANCOM-BC2 method. Complete results across all samples were presented in Supplementary Table 6.

The global analysis identified seven taxa whose relative abundances showed significant variation with lake size through a mass and surface area interaction. We fitted lake-specific models and extracted per-lake log fold change estimates for mass on these taxa. Lake-specific statistical results were summarized in Supplementary Table 7.

Differential abundance analysis showed that the relationship between fish mass and microbial taxa abundance was strongly dependent on lake size. The log fold changes of all selected taxa with fish mass exhibited an inverted-U shaped pattern, indicating that taxa in intermediate-sized lakes tended to show positive associations between their relative abundance and fish mass, whereas those in both small and large lakes tended to show weaker or negative associations (Figure 5A). The relative abundance of *Terrimicrobium* and *uncultured_f_Anaerolineaceae* showed linearly positive associations with fish mass and lake size (Figure 5B). *Rickettsia* and *uncultured_f_Isosphaeraceae* exhibited nonlinear patterns, showing strong negative associations with fish mass in small lakes that weakened in intermediate- and large-sized lakes. In contrast, *Hyphomicrobium* showed the opposite trend, shifting from positive associations in small lakes to weakly negative associations in intermediate and large lakes. The relative abundance of *ZOR0006* displayed an inverted-U shaped relationship, with the strongest positive association between fish mass and relative abundance in intermediate-sized lakes (Figure 5B). Overall, these findings demonstrate that mass microbiota relationships are lake-size dependent, with the strongest and most consistent signals emerging in the largest lakes, where significant associations were predominantly negative.

**Figure 5.**
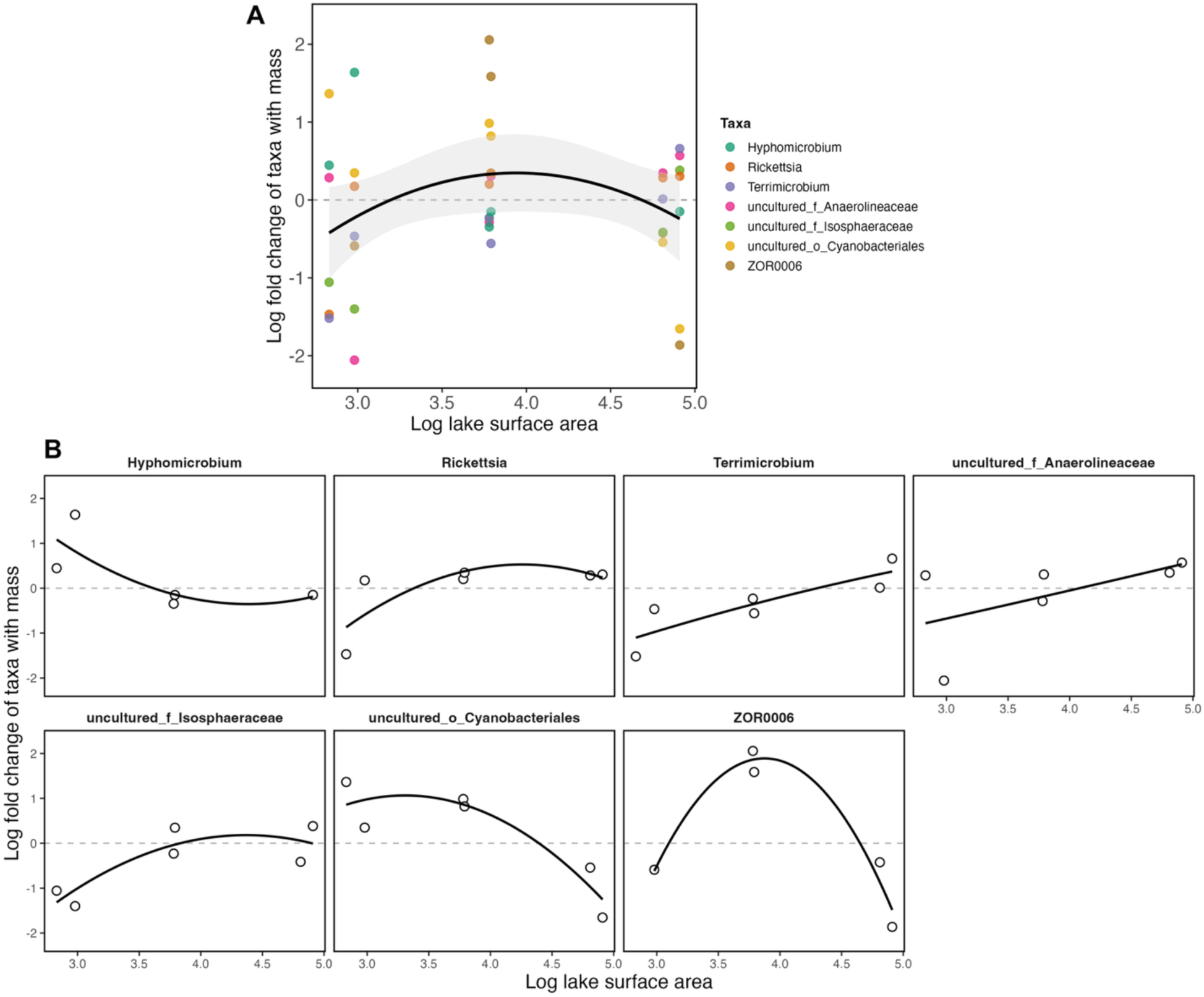
Differential abundance analysis showed fish mass influenced microbial taxa differently across lake sizes using ANCOM-BC2. A total of 7 taxa showed significant association with the interaction between fish mass and log lake surface area in the global analysis and were selected for lake-specific models. For each lake, log fold change in taxa abundance with fish mass was shown in relation to its log-transformed surface area. Taxa absent from a given lake are not shown as dots. Curves show fitted trends for visualization. Statistical results of taxa in each lake were shown in Supplementary Table 7.

## Discussion

This study examined how environmental (lake surface area, ecotype) and biological (sex, gravidity, *S. solidus* infection, fibrosis score, mass) factors influence the gut microbiota of wild threespine stickleback across six Alaskan lake populations. We found that lake surface area significantly explained fish gut microbial community diversity, with intermediate-sized lakes exhibiting higher Shannon diversity than those from small and large lakes. However, this effect was not explained by benthic and limnetic ecotypes. Body mass was negatively correlated with Chao1 richness. The strengths and directions of mass and gut microbial diversity relationships varied with lake size, being weakest in intermediate lakes and stronger in both small and large lakes.

In the current study, lake surface area had a quadratic effect on stickleback gut microbiota alpha diversity, with intermediate-sized lakes exhibiting greater alpha diversity than small or large lakes. One plausible mechanism is that dietary heterogeneity varies with lake surface area. It is possible that stickleback populations consumed more balanced proportions of benthic and limnetic prey and displayed the highest among-individual diet variation in intermediate-sized lakes [13]. Although we did not directly measure diet in this study, evidence from fish and other systems suggests that the consequences of diet diversity for gut microbiota are mixed. For example, Bolnick et al. found that in freshwater fish (wild-caught and lab-raised threespine stickleback, and Eurasian perch), individuals classified as dietary generalists (a mix of benthic and limnetic prey) often had lower gut microbial diversity than dietary specialists, with this effect being sex- and size-dependent in perch but independent of sex in stickleback [42]. By contrast, human studies showed a positive association between dietary diversity and alpha diversity of the gut microbiota [43]. Greater gut microbiota uniqueness was found in stickleback individuals from intermediate-sized lakes, indicating that increased diet diversity may promote gut microbiota diversity [14].

Because diet and trophic niche are closely tied to ecotype in stickleback, variation in feeding behavior and habitat use may also influence microbiota composition through ecomorphological differentiation [10, 44]. However, we found differences in lake surface area influencing gut microbiota composition were independent of benthic and limnetic ecotypes. It is important to note that our sampling occurred in spring, when both ecotypes tend to occupy the littoral zones and feed on similar macroinvertebrate resources. For example, in Spirit Lake (classified as limnetic), we observed many stickleback fish feeding on macroinvertebrates (benthic prey). By summer, as benthic resources became depleted and stickleback moved out from their nests, limnetic ecotypes typically shifted toward zooplankton feeding (limnetic prey) in the pelagic zones of deeper lakes. Therefore, dietary overlap early in the spring season may have reduced ecotype differences in gut microbiota composition in this study.

Beyond this seasonal context, a broader pattern is that ecotype composition often covaries with lake size. It is important to note that large and deep lakes tend to harbor more limnetic ecotypes, while small and shallow lakes tend to be more benthic [4]. However, this general pattern was not observed in the Alaskan lakes analyzed in this study. For example, the largest lake (Finger Lake, depth=13.4 m) harbored more benthic fish aligning with its extensive littoral habitat, while the smallest lake (Long Lake, depth=7.9 m) was dominated by limnetic fish and is characterized by a very hard bottom substrate that is unfavorable for macroinvertebrate development [27]. This deviation likely reflects the limited number of lakes examined and the inclusion of a few exceptional cases, rather than a true absence of correlation, and thus might not contradict broader regional trends. As a future direction, stable isotope analyses of stickleback muscle tissue could provide a more accurate assessment of individual dietary habits. Such methods integrate dietary intake over extended periods and could help quantify the proportions of benthic and limnetic prey consumed, thereby offering a more precise indication of ecotype contributions to gut microbial variation [45].

Our analyses revealed that body mass was a strong predictor of stickleback gut microbiota, and its effects were context dependent. We found that mass was negatively correlated with gut microbiota alpha diversity. This relationship could potentially be explained by several mechanisms. Smaller fish may exhibit greater metabolic rates than larger fish, influencing gut microbiota composition through altered behaviors or diet selection [9, 46–49]. Additionally, smaller adult fish may occupy specific ecological niches or microhabitats, such as different water currents or food sources, which expose them to distinct microbial communities compared to larger individuals [50]. All individuals sampled in the current study were adults, thereby eliminating the confounding effects of the dynamic gut microbiota changes typically observed during early developmental stages [51–53]. In addition, the strength and direction of this mass-gut microbiota relationship varied with lake surface area: negative in small lakes, weak in intermediate lakes, and positive in large lakes. One likely reason is that the effect of mass on gut microbiota is mediated by ecological context rather than absolute size differences since body mass distributions were comparable across lakes. In small lakes with limited habitat complexity and fewer prey types, larger fish may be forced to focus on the dominant prey and use more uniform microhabitats, thereby reducing the variety of microbial sources they encounter. Thus, increasing body mass in these settings may lead to a decrease in microbial richness. In large lakes, although limnetic prey often dominate, larger fish might access underexploited benthic zones, peripheral microhabitats, or rare prey types (e.g. nearshore, deep invertebrates), which expands their microbial exposure beyond diet. In intermediate lakes, where benthic and limnetic resources are more evenly mixed [42] and overlap among fish is higher, increasing mass may not lead a fish to new prey or habitats, resulting in minimal shifts in microbial exposure. The results of this study highlight that body mass affects gut microbiota in a context-dependent manner, varying across lake sizes, and underscore the importance of considering ecological settings when interpreting host-microbe associations in wild populations.

Our study offers new insights into how environmental and host traits jointly shape microbial communities. However, a key limitation is that the study is cross-sectional, limiting our ability to assess temporal dynamics in gut microbial communities. Future research should incorporate isotope analyses to more accurately distinguish benthic and limnetic ecotypes based on diet, as well as longitudinal sampling across different time points to determine whether the observed gut microbiota patterns are consistent over time and responsive to changing ecological conditions.

## Conclusion

In summary, this study offers a comprehensive understanding of how environmental factors and host traits impact gut microbiota composition in wild Alaskan threespine stickleback populations. Lake surface area was a strong ecological predictor, with intermediate-sized lakes supporting the highest microbial diversity. Body mass also influenced gut microbiota, but its effects were context-dependent, varying with lake size. These findings highlight that host-microbe interactions in stickleback are potentially mediated by ecological setting, underscoring the importance of integrating environmental and host-level perspectives to understand the eco-evolutionary dynamics of gut microbial communities in natural populations.

## Data availability statement

Sequence data have been deposited in the Sequence Read Database (SRA) under BioProjectID PRJNA1363120. The datasets analyzed in the current study can be accessed at https://github.com/sihanbu/Lake-size-and-stickleback-gut-microbiota

## Acknowledgement

This study was conducted with the support of the Alaska Department of Fish and Game (ADFG). The field work was carried out with help from Christopher Peterson, Kristofer Sasser, Elsa Diffo Tiyao, and Racheal Kramp. We are especially grateful for the foundational work conducted by our collaborators and landowners, which made this study possible.

