## Supplementary Figures for "Lake size shapes the relationship between body mass and gut microbiota in threespine stickleback (*Gasterosteus aculeatus*)"

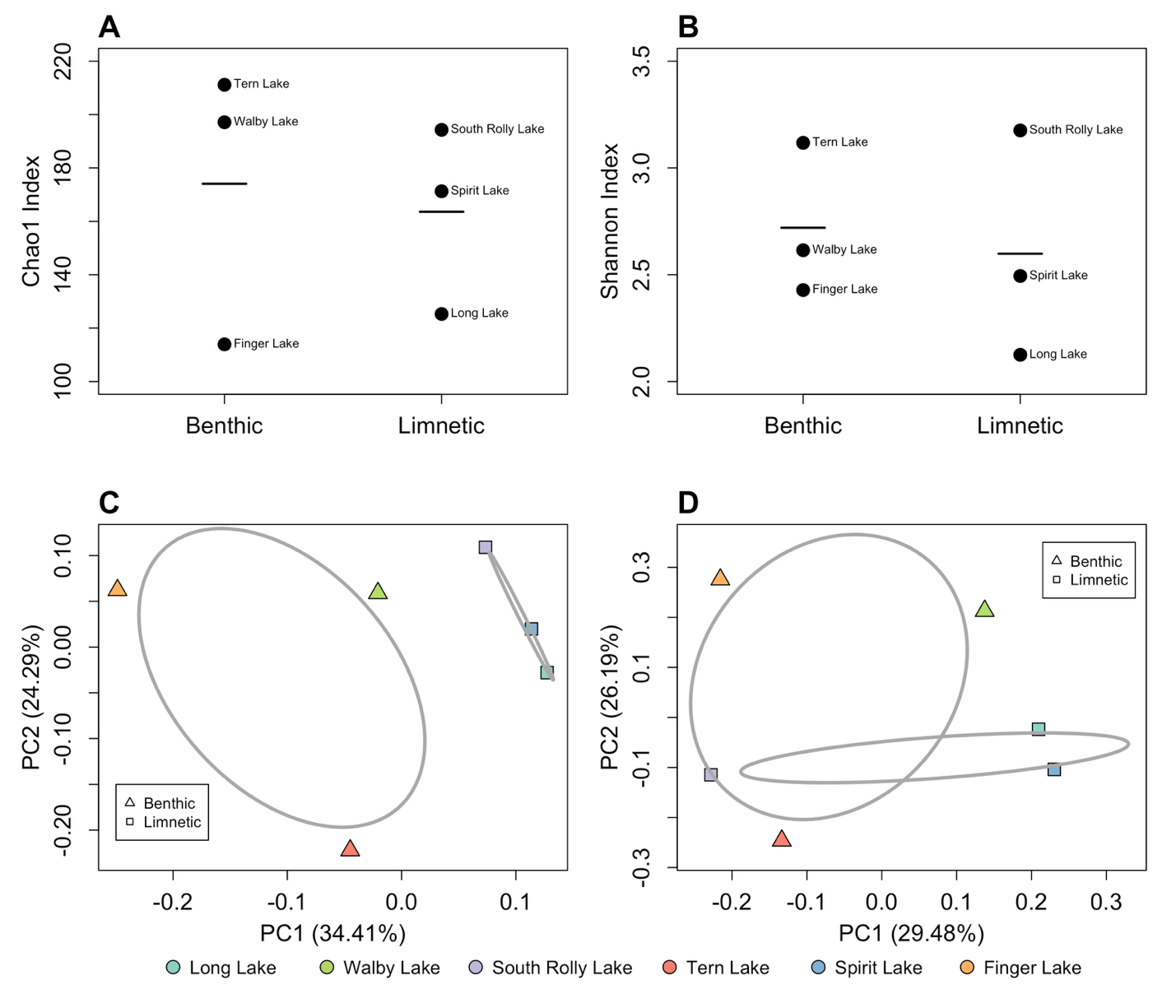

**Supplementary Figure 1.** Ecotype did not influence the fish gut microbiota alpha and beta diversity. The Wilcoxon rank-sum exact test showed no significant difference in (A) Chao1 index (*p*-value=0.7) and (B) Shannon diversity between benthic and limnetic lakes (*p*-value=1). Black lines represent the mean values for each lake mean. Similarly, no significant differences were detected for either the (C) Sorensen (F=1.58, R^2^=0.28, *p*-value=0.1) or (D) Bray-Curtis index (F=1.19, R^2^=0.23 *p*-value=0.3) of beta diversity using PERMANOVA test. The small sample size (n = 6) limited permutation counts and statistical power, making p-values coarse. Despite clear group separation in the ordination plot, significance was not detected. Ellipses indicate 95% confidence intervals around centroids.

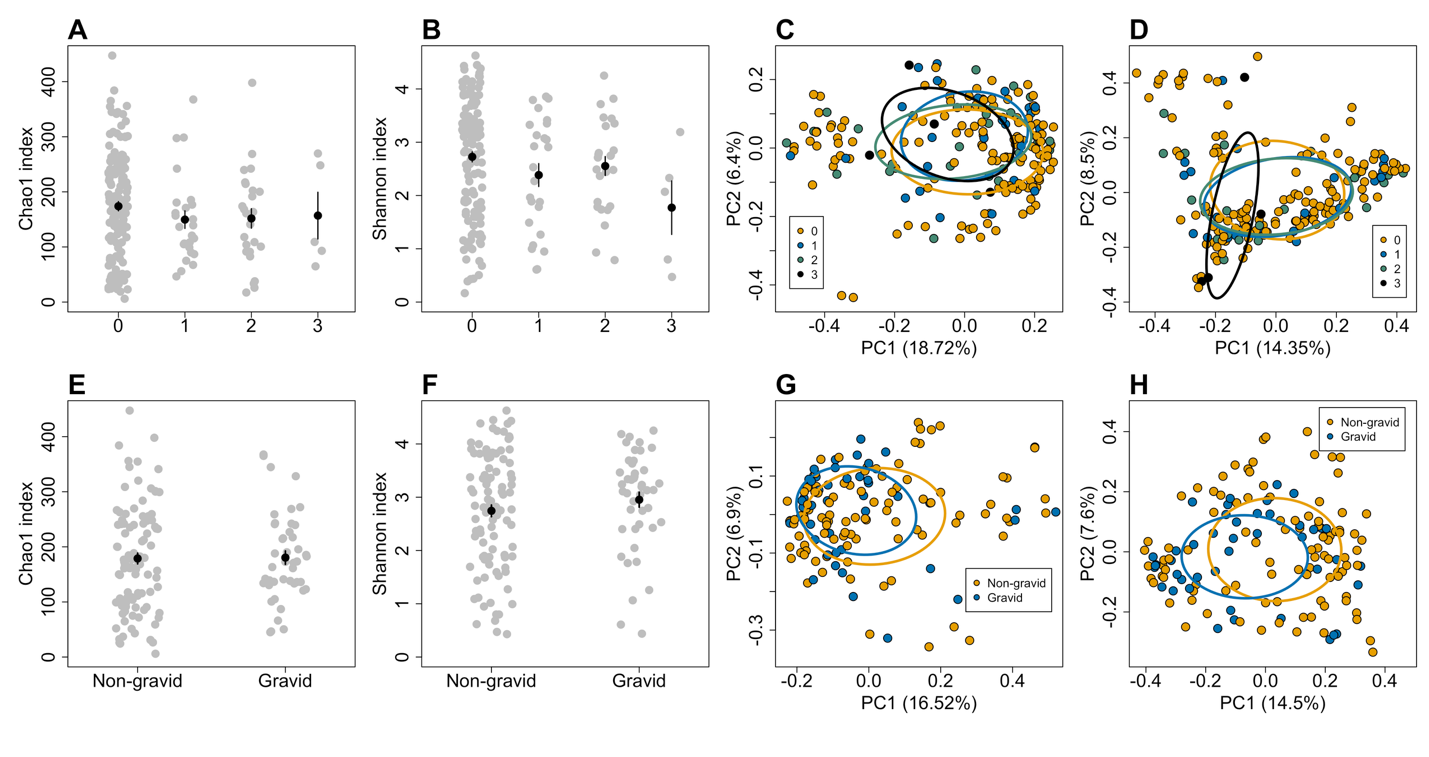

**Supplementary Figure 2**. Associations of fish gravidity and fibrosis with gut microbiota. All-lake analysis showed that fibrosis infection did not affect fish (A) Chao1 (F=0.34, *p*-value=0.8) and (B) Shannon index (F=1.35, *p*-value=0.26) of alpha diversity, and (C) Sorensen (F=0.96, R^2^=0.014, *p*-value=0.52) and (D) Bray-Curtis matrix (F=1.22, R^2^=0.02, *p*-value=0.11) of beta diversity. Among female fish, gravidity did not influence fish gut microbiota alpha diversity of (E) Chao1 (F=0.34, *p*-value=0.56) and (F) Shannon index (F=2.12, *p*-value=0.15) nor did it affect the beta diversity of (G) Sorensen dissimilarity matrix (F=1.27, R^2^=0.008, *p*-value=0.14) and (H) Bray-Curtis (F=1.46, R^2^=0.009, *p*-value=0.07). Black dots indicate the mean, and error bars are the mean±SE.
